# One Number to Rule Them All: The Wildlife Sperm Index for Standardized Gamete Assessment

**DOI:** 10.1101/2025.08.13.670140

**Authors:** Leah Jacobs, Andrew Benson, Joshua Jacobs, Kira L Marshall, Natalie E Calatayud

## Abstract

In wildlife conservation, breeding programs focused on reintroduction are critical to recovering endangered species. Assisted reproductive technologies (ARTs) and biobanking play pivotal roles in these efforts but depend on high-quality gametes. Spermatozoa, due to their structural simplicity and well documented quality assessments are often considered the preferred cell type for cryopreservation and ARTs, especially in rare or exotic species. However, inconsistent retrieval of sperm samples with sufficient volume and concentration, especially for critically endangered species, can pose major limitations to assessing overall sperm quality. To address this information gap, we have developed the Wildlife Sperm Index (WSI), a standardized scoring system that integrates key sperm quality metrics—such as motility, viability, and DNA integrity—into a weighted composite score. While this study applies the WSI to sperm, the framework is adaptable to other cell types used in biobanking and reproductive technologies, broadening its potential utility. Unlike traditional sperm assessments that rely solely on microscopy, the WSI provides a scalable and quantitative framework for evaluating samples across species, including those that yield low-volume or concentration sperm samples. We validated the WSI against traditional means of sperm quality assessments, demonstrating its reliability as both a quality assessment and decision-making tool, enabling users to evaluate sample suitability for cryopreservation, ARTs, or downstream applications. By enabling standardized quality tracking, the WSI provides a foundation for more informed reproductive decisions to support global conservation efforts.

**Summary sentence:** We present the Wildlife Sperm Index (WSI), a standardized, composite scoring system that integrates key sperm quality metrics to improve gamete assessment and supports assisted reproduction and biobanking in wildlife conservation.

## Introduction

The global conservation community is increasingly prioritizing the collection and preservation of viable cellular resources, including gametes, embryos, somatic cells, and cultured cell lines, to safeguard biodiversity and strengthen genetic management capacity for threatened and endangered species. The functional quality of these cells at the time of collection is a key determinant of their suitability for a wide range of applications, including assisted reproductive technologies (ARTs), ex situ research, genetic analyses, and long-term biobanking. Conservation breeding programs (CBPs) aimed at population reintroductions depend heavily on access to viable gametes for reproductive interventions such as hormone-assisted breeding, artificial insemination, or in vitro fertilization, as well as for downstream cryopreservation [1–3].

Maintaining high gamete quality standards at the point of collection is essential to ensure both immediate and long-term viability across diverse conservation applications. Poor-quality cells are more vulnerable to functional impairment, damage during processing or freezing, and reduced performance in reproductive procedures or laboratory assays [4–7]. These challenges are particularly acute for threatened species, where each available sample represents irreplaceable genetic diversity critical for sustaining shrinking populations.

While the principles of cell viability and quality assessment apply broadly across many cell types used in conservation, spermatozoa represent one of the most widely utilized and well- characterized cell types in wildlife biobanking and assisted reproduction. In addition, the availability of existing quality assessment methods, combined with their extensive use across taxa, makes sperm an ideal model for developing and validating standardized quality indexing frameworks [5, 8–14].

To address the need for standardized assessments, this project aims to develop a Wildlife Sperm Index (WSI), dedicated to evaluating spermatozoa quality. The goal is to provide a standardized methodology for assessing baseline values for cells obtained from wildlife species. Unlike the plentiful data available for humans and domestic animals, baseline parameters for wildlife are not often described or lack consistency in variables examined [5, 6, 14, 15]. As in humans and domestics, metrics like concentration and motility, acrosome and mitochondrial integrity, DNA degradation, and other physiologically relevant factors such as pH and osmolality are crucial for wildlife sperm quality [5, 8, 11, 13, 14, 16, 17].

To tackle both the variability in available parameters and the species-specific differences in their relative importance, the WSI incorporates multiple quality metrics into a scoring system using weighted averages. The WSI provides detailed viability scores, supporting decision-making in reproductive technologies and enabling biobanks to maintain a standardized sperm quality evaluation method [5, 18]. The WSI’s significance lies in its potential to revolutionize global conservation practices as a standardized framework for sperm quality assessment. It functions as an open-access quality indexing tool, particularly beneficial for rare or limited-volume samples, allowing researchers to catalog, standardize, and track sperm sample quality throughout collection and storage workflows [1, 5, 11, 16, 19]. In addition to standardizing sperm quality assessments, this technique provides a distinct advantage by enabling more informed decisions about the use of sperm samples. For species where high-quality samples are scarce or re- collection is impractical, the index can guide practitioners in adapting handling protocols or selecting fertilization methods, such as ICSI, to maximize the reproductive potential of all samples, not just those of optimal quality. This is particularly important in conservation and captive breeding programs, where maximizing the utility of every sample is critical for preserving genetic diversity. As conservation breeding efforts expand, they face increasing logistical and financial challenges, underscoring the importance of ARTs and biobanking as essential tools for genetic management [20]. Tools like the Wildlife Sperm Index enhance gamete quality assessments, supporting breeding programs and reproductive strategies vital for conserving endangered species and maintaining biodiversity.

## Methods

### Mathematical Models

#### Data Standardization

Grading sample quality is essential for maintaining unbiased and consistent metrics in sample storage and future applications. In this study, we implemented a Weighted Average filter to evaluate complex data sets with multiple measurements, assigning each sample a singular quality value (Fig. 1).

**Figure 1.**
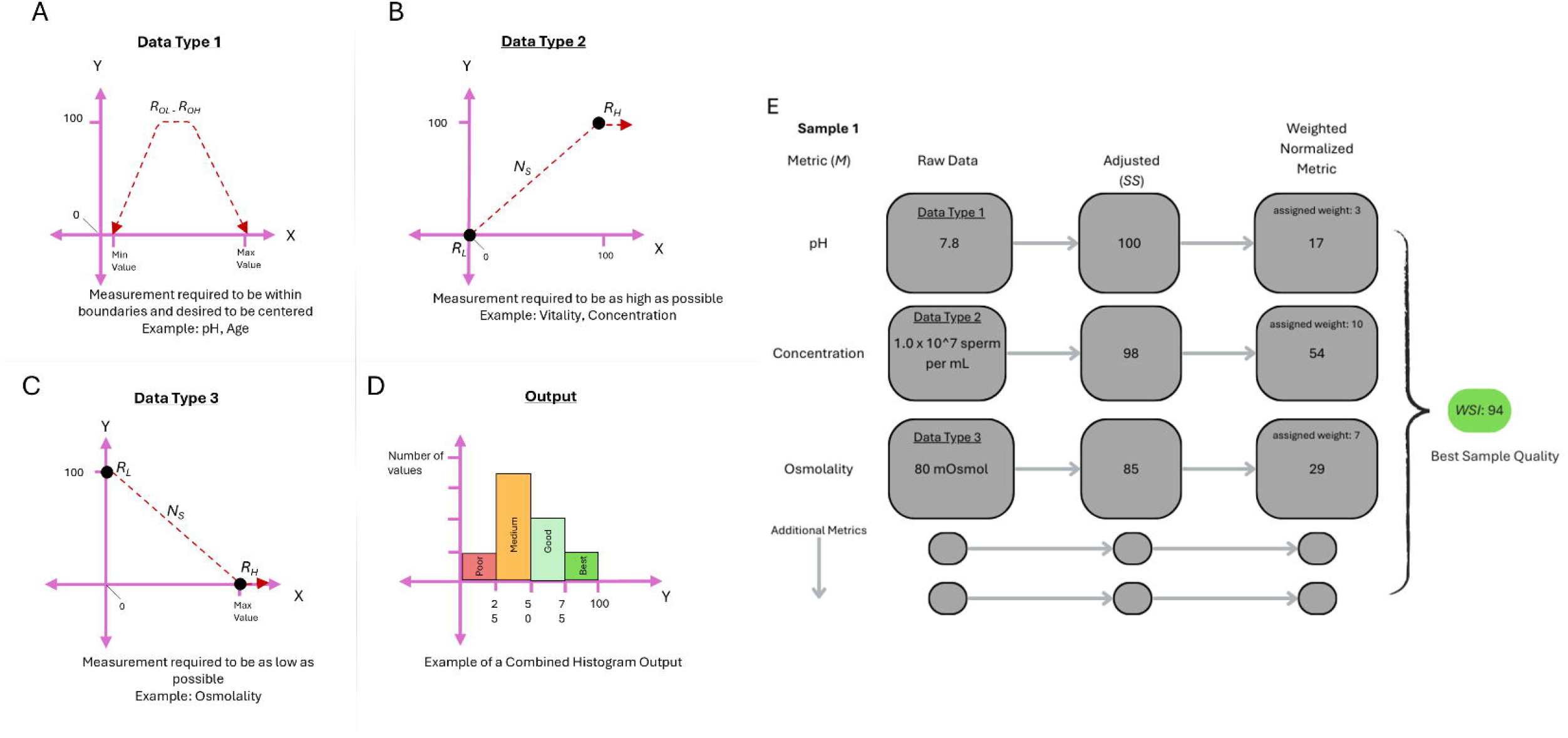
Data adjustment and binning. 1A. Data normalization for metrics with optimal range within a specific set of boundaries with either extreme indicating a poor quality metric (i.e. for pH). 1B. Data manipulation for data on a linear scale, where the greater the value, the better the quality (i.e. Concentration, Volume). 1C. Data manipulation for data on an inverse linear scale, where the lower the value, the optimal the sample (i.e. Osmolality). 1D. Histogram highlighting potential distribution of samples in a dataset. 1E. Flow chart of creating the index starting with adjusting the data to a 0-100 unitless scale, assigning a weight and generating an index value combining all metrics available.

The Weighted Average filter comprises four key components: Samples, Metrics, Weights, and Range. **Samples** refer to the physical specimens selected for evaluation. **Metrics** are the specific attributes used to assess the quality of these samples (e.g. volume and pH). Weights are assigned to each metric to reflect the relative importance of these attributes; for example, volume may be weighted twice as heavily as pH when determining overall sample quality. **Range** defines the acceptable values for each metric, establishing criteria such as a preferred pH range of 7.6-8.2 (see Table 1).

**Table 1.**
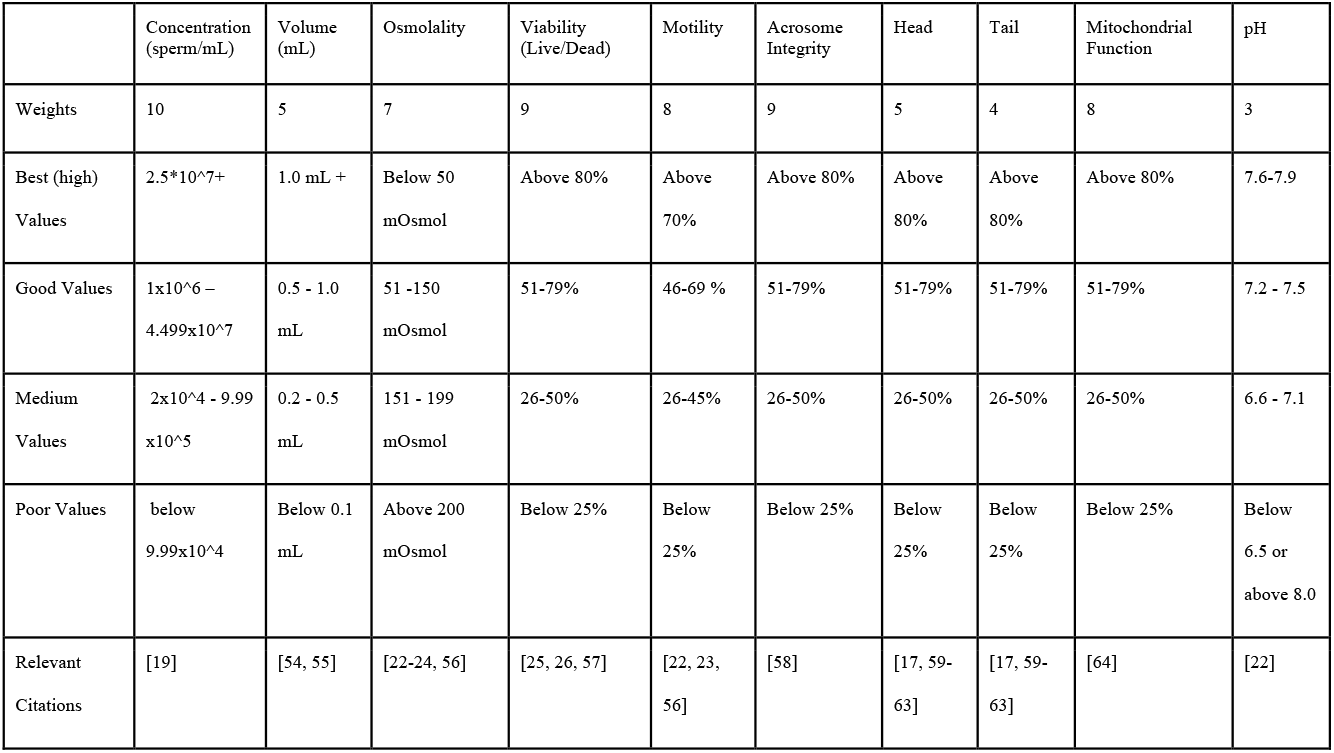
Ranges for each metric are assessed based on values obtained from peer-reviewed literature on *Xenopus laevis* and other anuran amphibians.

|  | Concentration<br>(sperm/mL) | Volume<br>(mL) | Osmolality | Viability<br>(Live/Dead) | Motility | Acrosome<br>Integrity | Head | Tail | Mitochondrial<br>Function | pH |
| --- | --- | --- | --- | --- | --- | --- | --- | --- | --- | --- |
| Weights | 10 | 5 | 7 | 9 | 8 | 9 | 5 | 4 | 8 | 3 |
| Best (high)<br>Values | 2.5*10<sup>7</sup>+ | 1.0 mL + | Below 50<br>mOsmol | Above 80% | Above<br>70% | Above 80% | Above<br>80% | Above<br>80% | Above 80% | 7.6-7.9 |
| Good Values | 1x10<sup>6</sup> –<br>4.499x10<sup>7</sup> | 0.5 - 1.0<br>mL | 51 -150<br>mOsmol | 51-79% | 46-69 % | 51-79% | 51-79% | 51-79% | 51-79% | 7.2 - 7.5 |
| Medium<br>Values | 2x10<sup>4</sup> - 9.99<br>x10<sup>5</sup> | 0.2 - 0.5<br>mL | 151 - 199<br>mOsmol | 26-50% | 26-45% | 26-50% | 26-50% | 26-50% | 26-50% | 6.6 - 7.1 |
| Poor Values | below<br>9.99x10<sup>4</sup> | Below 0.1<br>mL | Above 200<br>mOsmol | Below 25% | Below<br>25% | Below 25% | Below<br>25% | Below<br>25% | Below 25% | Below<br>6.5 or<br>above 8.0 |
| Relevant<br>Citations | [19] | [54, 55] | [22-24, 56] | [25, 26, 57] | [22, 23,<br>56] | [58] | [17, 59-63] | [17, 59-63] | [64] | [22] |

To standardize all Metrics on a unitless scale, data are normalized to a 0 - 100 range. This process was carried out by mapping each of the metrics to a standardized, unitless scale from 0 - 100, with 100 indicating the best possible value for that metric, and zero reflecting the worst possible value. Each metric was adjusted based on their optimal values (standardized per species or group of species) and was categorized into one of three data types. Data type 1 includes data that falls within an optimal medium range and is suboptimal or poor as values reach each extreme. Age range is an example of data type 1 where younger and older ages tend to be less reproductively viable. Data type 2 metrics follow a linear scale where 0 represents the lowest performance, while 100 is the highest. Metrics in this data type include concentration, volume, live/dead, and motility. Data type 3 metrics, in contrast, follow an inverse linear scale where values approaching 100 are considered suboptimal and lower values are considered optimal.

Osmolality, for instance, functions as a type 3 metric in many freshwater aquatic species, where lower osmolality indicates active, viable sperm, while high osmolality results in inactivity and lower fertilization potential.

Each data type requires a specific normalization process to map on the unitless scale.

Data type 1:

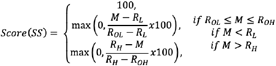

Data type 2:

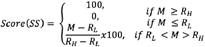

Data type 3:

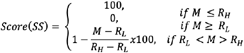

Where: *R*_*H*_: Range High; *R*_*L*_: Range Low; *R*_*OL*_: Range Optimal, Low end; *R*_*OH*_: Range Optimal, High end; M: Metric value; SS: Adjusted Sample Score

This normalization step ensures that all metrics contribute proportionally to the final index, independent of their original units.

#### Weighted Averages

To calculate the weighted average and develop the WSI, the model incorporates all applicable metrics and calculates a single value based on the weight assigned to each metric. Weighting factors are determined based on the relative significance of each metric for each species using species knowledge, literature reviews, expert opinion, or experimental validation.

Calculated WSI:

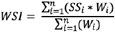

Where: W: Weight; SS: Adjusted Sample Score

### Graphical User Interface Development

To maximize the accessibility and usability of the Wildlife Sperm Index for a variety of contexts and across institutions, we designed a graphical user interface (GUI) using python 3.14 (Fig. 2). This application creates a platform that can be utilized by anyone to assess multiple characteristics of sperm quality and develop an index. We highlight the components of the GUI below:

**Figure 2.**
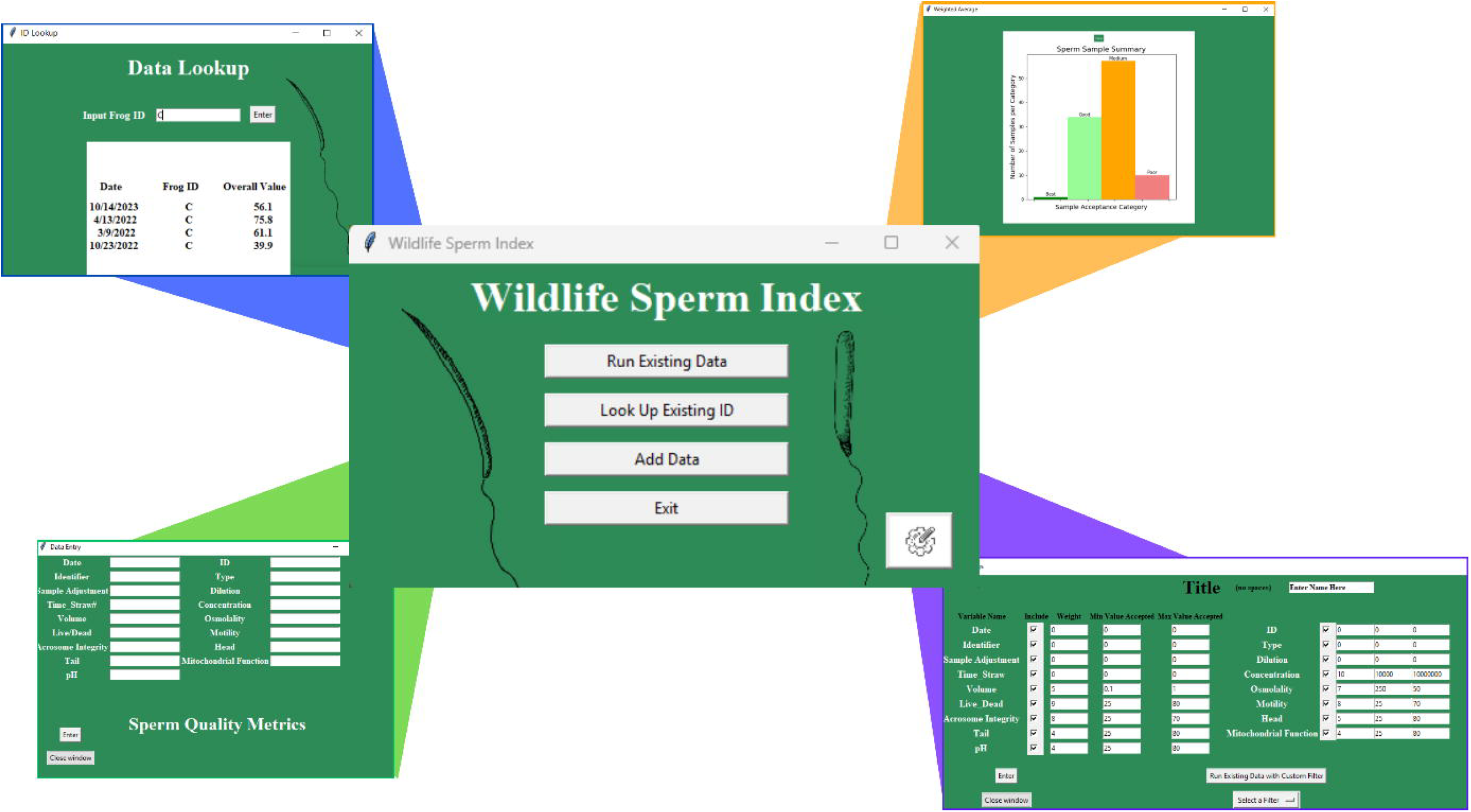
Wildlife Sperm Index GUI with main terminal and outputs for each button including the ‘Run existing Data’ button which calculates the sperm index and summarizes the data (top right), ‘Look Up Existing ID’ button that allows for individual specimen examination over time (top left), ‘Add Data’ button that allows for real time addition of data (bottom left), and the settings button (bottom right) that allows for the adjustment of the metric parameters including assigned weights and low and high values.

#### GUI main page

The main terminal has five buttons which are described in more detail below: “Run Existing Data”, “Look Up Existing ID”, “Add Data”, a settings button, and “Exit”.

#### Data Import and Scoring

The “Run Existing Data” functionality was developed to import an Excel file pre-populated with metrics and samples to evaluate adjusted sample score (SS). Upon execution, the system calculates the SS and appends a metric adjustment (MN) score to the Excel file, allowing for further investigation into the factors affecting the scores. The script calculates the sperm index using the same formulas as described above. All values are assessed using the weights assigned in the settings tab. The index is populated into a column within the specified Excel sheet called “Overall Score.” Additionally, a graphical scatter plot is generated to visually represent all samples and corresponding SS values, providing insight into batch trends and the precision of testing methodologies (Fig. 2).

The system features alternative interface capabilities, including the “Look Up Existing ID” function, which queries a specified Excel file for reported sample scores to evaluate their usability across time or treatment.

The “Add Data” function assesses all metrics in the Excel file and presents input boxes for data collection; upon providing the requested information, the Excel file updates with the new values.

Additionally, the settings tab permits the execution of an alternative algorithm, allowing users to modify previously fixed parameters such as the upper and lower limits for their metrics (based on the species assessed), and weightings (W) for each metric. This functionality generates alternative SS values while preserving the original SS, which is essential for repeatable sample comparison. These alternative SS values are beneficial for evaluating different species, accommodating changes in allowed ranges or conducting hypothetical analyses.

### Case study - Aquatic amphibian sperm quality

To validate the efficacy of the WSI and assess the usability of the GUI, a set of data was created based on the well-documented sperm quality parameters of two different aquatic amphibian species: the African clawed frog, a well-documented model organism (*Xenopus laevis*) and the Mountain yellow-legged frog, a critically endangered amphibian from Southern California (*Rana muscosa*). Metrics and weights were assigned based on literature review and the authors knowledge of aquatic amphibian sperm (Table 1 & 2).

#### Testing the index using a Model species

As a proof of concept, a specific set of formulated values was created to assess the applicability of the scoring index based on well documented sperm metrics for the African clawed frog [19, 21–28]. For the first run through, we wanted to examine as many metrics as possible. The metrics examined included pH, volume, motility, osmolality, acrosome integrity, mitochondrial function, basic morphology (head and tail), viability, and concentration. Each metric was assigned values that reflected the biological relevance of the metric for this species. Samples first underwent manual evaluations where each metric was examined, and corresponding values were assigned manually by using Excel calculations before being processed through the GUI to confirm the usability of the program. Central to the WSI framework is its normalization process, which converts all metrics to a unitless 0–100 scale, enabling proportional comparisons across diverse measurements. This normalization ensures that all included metrics contribute appropriately to the final weighted score, regardless of their original scale or units. The final WSI score is calculated as a weighted average of the adjusted sample scores for each available metric. Weights are assigned based on species knowledge, empirical evidence, or expert judgment, allowing the model to emphasize biologically informative metrics while de-emphasizing those less relevant or more variable. Crucially, when certain metrics are missing for a given sample, the WSI calculation automatically excludes these from the weighted sum without penalizing the final score. This adaptive feature allows for consistent evaluation even when complete datasets are not attainable—an inherent limitation of working with rare or limited wildlife samples.

## Results

For any cell left blank (i.e. the metric was not recorded in quality assessments) the index ignores the metric and calculates the weighted average based on the available data. Intra-class correlation was high between GUI calculated indices and Excel calculated indices, confirming the ease of use and applicability of the Wildlife Sperm Index application. The intra-class correlation coefficient (ICC) between WSI and Qualitative scores was 0.99, indicating excellent agreement between the two methods (p < 0.001, 95% CI: 0.985–0.993). These results suggest the methods are highly consistent and may be used interchangeably in this context.

To further test usage of the GUI using a ‘real-world’ application, a set of data collected from spermiation trials for the Mountain yellow-legged frog was used. This species is a highly endangered southern California native amphibian whose reproduction in human care is inconsistent and difficult to achieve [29]. Reproductive technologies are utilized to help increase reproductive output, including the use of exogenous hormones to induce spermiation and ovulation. Protocols have been developed for the collection of sperm from this species, however, sperm output and volume is generally very low and, being an endangered species, irreplaceable. Because of the low output of volume given by this species during hormone induced spermiation techniques, only a handful of metrics are assessed for each sample. Typically, we are restricted to assessing concentration, motility, live/dead and volume obtained. Metric parameters were chosen based on the authors knowledge of the quality of sperm in this species (Table 2). All metrics were changed in the ‘settings’ feature within the GUI to test the ease of species-specific parameter adjustments. As with the example above, data was first assessed via Excel calculations prior to being processed with the GUI. Intra-class correlation was high between GUI calculated indices and Excel calculated indices, confirming the ease of use and applicability of the Wildlife Sperm Index application. The intra-class correlation coefficient (ICC) between WSI and Qualitative scores was 1, indicating perfect agreement between the two methods (p < 0.001, 95% CI: 1-1). This dataset was a much smaller dataset and reflective of the typical quality parameters available from *Rana muscosa*. Because of this, qualitative values were easily assessed and given based on our knowledge, reflecting the values also generated by the WSI. These results suggest the methods are highly consistent and may be used interchangeably in this context, especially when practitioners have such in-depth knowledge of their species and the parameters available.

**Table 2.** Ranges for each metric are assessed based on values obtained from author knowledge for the Mountain yellow legged frog (*Rana muscosa*). Calatayud et al. 2025 (in prep.).

|  | Concentration<br>(sperm/mL) | Volume<br>(mL) | Osmolality | Viability<br>(Live/Dead) | Motility | Acrosome<br>Integrity | Head | Tail | Mitochondrial<br>Function | pH |
| --- | --- | --- | --- | --- | --- | --- | --- | --- | --- | --- |
| Weights | 10 | 9 | 7 | 6 | 2 | 9 | 5 | 4 | 8 | 3 |
| Best<br>(high)<br>Values | 7.0*10<sup>6</sup>+ | 600 uL<br>+ | Below 75<br>mOsmol | Above 80% | Above<br>80% | Above<br>80% | Above<br>80% | Above<br>80% | Above 80% | 7.0-7.2 |
| Good<br>Values | 1.0 x10<sup>6</sup> –<br>6.99 x10<sup>6</sup> | 400-600<br>uL | 76 -100<br>mOsmol | 51-79% | 51-79% | 51-79% | 51-79% | 51-79% | 51-79% | 7.2 –<br>7.8,<br>6.0-7.2 |
| Mediu<br>m<br>Values | 2.0 x10<sup>4</sup> -<br>9.9 x10<sup>5</sup> | 200-400<br>uL | 101 - 200<br>mOsmol | 26-50% | 26-50% | 26-50% | 26-50% | 26-50% | 26-50% | 5.8-5.9,<br>7.9-8.1 |
| Poor<br>Values | below<br>1.99x10<sup>4</sup> | Below<br>100 uL | Above 201<br>mOsmol | Below 25% | Below<br>25% | Below<br>25% | Below<br>25% | Below<br>25% | Below 25% | Below<br>5.7 or<br>above<br>8.2 |

## Discussion

The Wildlife Sperm Index (WSI) provides a standardized, flexible framework for evaluating sperm and cell quality with a broad applicability across taxa [29–31]. By integrating multiple metrics the WSI allows researchers to generate a comprehensive, objective quality score even when only partial datasets are available [6, 8, 9, 15, 32]. Species-specific weightings and ranges can be assigned for each metric, reflecting biological relevance and allowing customization to the physiological characteristics of different taxa [13, 33].

Traditional sperm quality indices developed for humans or domestic animals often rely on high-volume ejaculates with commonly measured parameters (e.g. concentration, motility, morphology)[7, 34–37]. These models typically assume full datasets and fixed weighting schemes that are poorly suited to non-model or endangered taxa, where sample volumes may be minimal and standard parameters may not reliably predict fertility. Few existing models for zoological or wildlife species offer comparable flexibility, and those that do often lack the quantitative standardization necessary to assess diverse sample types or incomplete datasets.

The WSI directly addresses these limitations, offering particular value for conservation programs where gametes are scarce, irreplaceable, and variable in quality. For many threatened or endangered species, obtaining sufficient sperm samples for conventional quality assessments is a persistent challenge. Males may produce minimal ejaculate volumes (e.g. felids, [38]), excrete sperm primarily in urine (e.g. amphibians [39]), or yield only a few spermatozoa upon collection (e.g. naked mole rat and some felids [40–45]). Traditional sperm analyses often require large sample volumes or rely on multiple parameters (concentrations, motility, morphology) to infer fertility potential[7, 13]. These criteria are often impossible to fulfill with minute or degraded samples. The WSI can integrate sparse or incomplete data into a standardized scoring framework and utilizing researcher assigned weights. By weighting critical parameters based on research knowledge of species sperm characteristics, the WSI generates a quality score, even when only one or two metrics are measurable. This approach is particularly vital for conservation programs where every gamete is considered irreplaceable. By maximizing insights from minimal material, the WSI reduces the risk of discarding viable samples due to perceived inadequacy or limited means of assessment.

To validate the model, we applied the WSI to two aquatic amphibian species that illustrate these challenges. The African clawed frog (*Xenopus laevis*) provided a proof-of-concept dataset incorporating multiple metrics, while the critically endangered mountain yellow-legged frog (*Rana muscosa*) provided real-world data with restricted metrics, reflecting field conditions where only volume, concentration, motility, and viability could be routinely assessed. In both cases, species-specific weighting schemes were implemented using the WSI’s graphical user interface (GUI), which allows users to upload data, adjust metric ranges and weights, and automatically calculate final scores. Importantly, the GUI validated calculations against manually processed Excel data, yielding an intra-class correlation coefficient (ICC) of 0.99 (p < 0.001; 95% CI: 0.985–0.993), confirming the reliability and reproducibility of the automated scoring system.

Although we focused primarily on aquatic amphibian species, the WSI can be applied and modified for other taxa. To apply the index to other taxa, practitioner knowledge must be utilitized to adjust the weightings for each specific species examined. For example, mammalian sperm challenges range from microspermia (small volumes) to cryosensitivity. For example, for felids such as cheetahs, captive males often produce <20 µL of semen with high teratospermia[46]. The WSI can discount morphology (if uniformly poor) and emphasize motility and speed of progression as well as acrosomal status. For reptiles, such as the Galapagos tortoise (*Chelonoidis nigra*), where semen collection is rare, the WSI can standardize scores from single epididymal sperm samples post-mortem to determine whether cryopreservation is optimal for the sample. Additionally, in snakes and lizards, sperm storage in females necessitates examination of overall sperm morphology, including acrosome integrity, DNA integrity and damage (i.e. Chromatin dispersion tests) over volume due to the low volumetric output and importance of morphology[47, 48].

Additionally, in other samples that may be obtained post-mortem, such as testes macerates, parameters such as morphology and viability may be assessed over motility which may not be present in osmotically balanced samples such as within the testes[49].

The WSI stands to be an open-access application that will allow the rapid and customizable assessment of sperm quality by combining multiple, applicable metrics into one decisive value. Figure 3 illustrates a decision-making framework developed to support the Wildlife Sperm Index (WSI), a tool designed to evaluate and standardize sperm quality across species using multiple biologically relevant parameters. The parameters assessed quantitatively by the WSI can ultimately be categorized into one of four quality tiers: Best-Quality, Good-Quality, Medium- Quality, and Poor-Quality, where Best- and Good-Quality samples are prioritized for artificial fertilization or cryopreservation (biobanking), while Medium-Quality samples may be used with compensatory techniques or reevaluated after further processing. Poor-Quality samples are either processed further or discarded, depending on their salvageability or potential utility for data collection.

**Figure 3.**
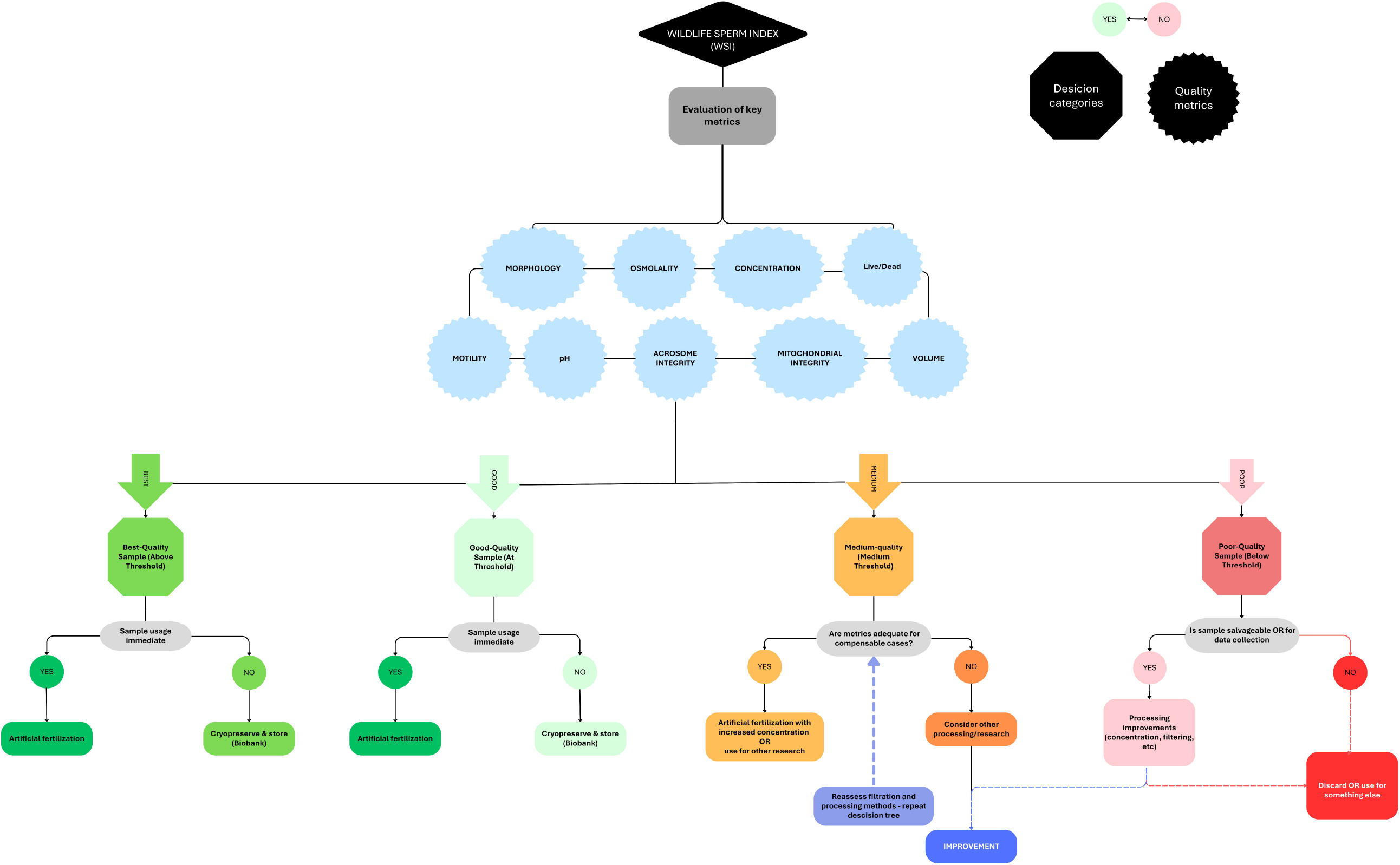
Decision tree evaluating sperm samples using the Wildlife Sperm Index (WSI). Samples are assessed for key quality metrics (e.g., motility, morphology, concentration) and categorized as high, medium, or low quality. High-quality samples are considered optimum and ready to be used for artificial fertilization or cryopreservation; medium-quality samples undergo further evaluation or processing; low-quality samples are either improved, repurposed, or discarded based on their potential for salvage or data use.

Finally, the WSI offers additional applications beyond individual sample assessment. Its flexible scoring platform can be used to compare outcomes across applications, such among storage techniques (e.g., fresh vs. frozen sperm), sperm processing techniques (spermic urine vs. macerates) and other factors affecting sperm quality. The integration of cryopreserved genetic data significantly enhances conservation efforts [50, 51], guiding breeding management to ensure optimal genetic diversity preservation under the One Plan Approach [52, 53]. Additionally, the WSI can facilitate comparisons in analytical techniques (e.g., flow cytometry vs. microscopy), aiding in the optimization of preservation and usage protocols [13, 33]).

By condensing multiple complex metrics into a single value, the WSI facilitates standardized decision-making, promotes optimal use of limited or sensitive genetic material, and improves conservation outcomes across diverse species.

This adaptability enhances its utility in conservation breeding programs (CBPs), cryopreservation efforts, and assisted reproductive technologies (ARTs), where maximizing the reproductive potential of every sample is critical for sustaining endangered populations. The WSI thus represents a substantial advance in wildlife reproductive science, addressing key analytical challenges in a highly practical, adaptable, and biologically grounded framework.

## Acknowledgments

The authors would like to acknowledge San Diego Zoo wildlife Alliance and the Mountain yellow-legged frog recovery team for their support and guidance regarding reproductive efforts and the use of a small dataset for mountain yellow-legged frog sperm characteristics.

## Author Contributions

LJ, AB, and JJ created the code and developed the graphical user interface (GUI) for the index; LJ and NC conceptualized the study, conducted background literature review, and drafted the manuscript; KM contributed to study conceptualization and manuscript editing; all authors contributed to writing and revising the manuscript and approved the final version for submission.

## References

1. De Mori, B., et al., The ethical assessment of Assisted Reproductive Technologies (ART) in wildlife conservation. Biological Conservation, 2024. 290: p. 110423.

2. Fujihara, M. and M. Inoue-Murayama, The wildlife biobanking of germ cells for in situ and ex situ conservation in Japan. Theriogenology Wild, 2024. 4: p. 100086.

3. Global Species Action Plan: Supporting implementation of the Kunming Montreal Global Biodiversity Framework. 2023, IUCN. Gland Switzerland.

4. Comizzoli, P., et al., Comparative cryobiological traits and requirements for gametes and gonadal tissues collected from wildlife species. Theriogenology, 2012. 78(8): p. 1666–1681.

5. Braundmeier, A.G. and D.J. Miller, Invited Review: The Search is on: Finding Accurate Molecular Markers of Male Fertility. Journal of Dairy Science, 2001. 84(9): p. 1915–1925.

6. Ezzati, M., et al., Influence of cryopreservation on structure and function of mammalian spermatozoa: an overview. Cell and Tissue Banking, 2020. 21(1): p. 1–15.

7. WHO laboratory manual for the examination and processing of human semen. 2021, World Health Organization.

8. Kowalski, R.K. and B.I. Cejko, Sperm quality in fish: Determinants and affecting factors. Theriogenology, 2019. 135: p. 94–108.

9. Quintero-Pérez, R.I., et al., Trade-off between thermal preference and sperm maturation in a montane lizard. Journal of Thermal Biology, 2023. 113: p. 103526.

10. Janosikova, M., et al., New approaches for long-term conservation of rooster spermatozoa. Poultry Science, 2023. 102(2): p. 102386.

11. Otero, Y., et al., Recovery and Characterization of Spermatozoa in a Neotropical, Terrestrial, Direct-Developing Riparian Frog (Craugastor evanesco) through Hormonal Stimulation. Animals, 2023. 13(17): p. 2689.

12. Arregui, L., et al., Hormonal induction of spermiation in a Eurasian bufonid (Epidalea calamita). Reproductive Biology and Endocrinology, 2019. 17(1).

13. van der Horst, G., Status of Sperm Functionality Assessment in Wildlife Species: From Fish to Primates. Animals, 2021. 11(6): p. 1491.

14. Aitken, R.J. and M.A. Baker, Oxidative stress, sperm survival and fertility control. Molecular and Cellular Endocrinology, 2006. 250(1-2): p. 66–69.

15. Canal Domenech, B. and C. Fricke, Recovery from heat-induced infertility—A study of reproductive tissue responses and fitness consequences in male <i>Drosophila melanogaster</i>. Ecology and Evolution, 2022. 12(12): p. e9563.

16. Hirohashi, N. and R. Yanagimachi, Sperm acrosome reaction: its site and role in fertilization†. Biology of Reproduction, 2018. 99(1): p. 127–133.

17. Arregui, L., et al., Determining the effects of sperm activation in anuran cloaca on motility and DNA integrity in. Reproduction, Fertility and Development, 2021. 34(5): p. 438–446.

18. Comizzoli, P. and W.V. Holt, Recent Progress in Spermatology Contributing to the Knowledge and Conservation of Rare and Endangered Species. Annual Review of Animal Biosciences, 2022. 10(1): p. 469–490.

19. Wolf, D.P. and J.L. Hedrick, A molecular approach to fertilization. Developmental Biology, 1971. 25(3): p. 348–359.

20. Engdawork, A., T. Belayhun, and T. Aseged, The Role of Reproductive Technologies and Cryopreservation of Genetic Materials in the Conservation of Animal Genetic Resources. Ecological Genetics and Genomics, 2024. 31: p. 100250.

21. Waggener, W.L. and E.J. Carroll, A method for hormonal induction of sperm release in anurans (eight species) and <i>in vitro</i> fertilization in <i>Lepidobatrachus</i> species. Development, Growth & Differentiation, 1998. 40(1): p. 19–25.

22. Christensen, J.R., et al., Effects of pH and dilution on African clawed frog (Xenopus laevis) sperm motility. Canadian journal of zoology, 2004. 82(4): p. 555–563.

23. Christensen, J.R., The effects of environmental contaminants on metamorphosis in Rana catesbeiana and sperm motility in Xenopus laevis. 2009.

24. Inoda, T. and M. Morisawa, Effect of osmolality on the initiation of sperm motility in Xenopus laevis. Comparative biochemistry and physiology. A, Comparative physiology, 1987. 88(3): p. 539–542.

25. Arregui, L., J.C. Koch, and T.R. Tiersch, Transitioning from a research protocol to a scalable applied pathway for Xenopus laevis sperm cryopreservation at a national stock center: The effect of cryoprotectants. Journal of Experimental Zoology Part B: Molecular and Developmental Evolution, 2024. 342(3): p. 291–300.

26. Morrow, S., et al., Effects of freezing and activation on membrane quality and DNA damage in Xenopus tropicalis and Xenopus laevis spermatozoa. Reproduction, Fertility and Development, 2016. 29: p. 1556–1566.

27. Ogawa, A., et al., Induction of ovulation in Xenopus without hCG injection: the effect of adding steroids into the aquatic environment. Reproductive Biology and Endocrinology, 2011. 9(1): p. 11.

28. Sargent, M.G. and T.J. Mohun, Cryopreservation of sperm ofXenopus laevis andXenopus tropicalis. genesis, 2005. 41(1): p. 41–46.

29. Calatayud, N.E., N. Gardner, and D.M. Shier, Captive breeding and reintroduction of the mountain yellow-legged frog (Rana muscosa). 2016 annual report, San Diego Zoo, Institute for Conservation Research Division of Applied Animal Ecology, Escondido, CA, 2016.

30. Takeshima, M. and A. Gotoh, Establishment of a rapid, cost-effective, and accurate method for assessing insect sperm viability. Journal of Insect Physiology, 2024. 158: p. 104682.

31. Bezerra, L.G.P., et al., Development of assays for the characterization of sperm motility parameters, viability, and membrane integrity in the epididymis and vas deferens of the greater rhea (Rhea americana). Animal Reproduction, 2023. 20(4): p. e20230113.

32. Jacobs, L., et al., Developing flow cytometry for precise evaluation of amphibian sperm viability: technical report. Reproduction, Fertility and Development, 2025. 37(4).

33. Tanga, B.M., et al., Semen evaluation: methodological advancements in sperm quality-specific fertility assessment - A review. (2765-0189 (Print)).

34. Chawre, S., et al., A Review of Semen Analysis: Updates From the WHO Sixth Edition Manual and Advances in Male Fertility Assessment. (2168-8184 (Print)).

35. Chenoweth, P., et al., Andrology laboratory review: evaluation of sperm morphology. Clinical Theriogenology, 2024. 16.

36. Chenoweth, P.J. and S.P. Lorton, Manual of animal andrology. 2022: CABI.

37. Hafez, E.S.E. and B. Hafez, Reproduction in farm animals. 2013: John Wiley & Sons.

38. Swanson, W., et al., Reproductive status of endemic felid species in Latin American zoos and implications for ex situ conservation. Zoo Biology: Published in affiliation with the American Zoo and Aquarium Association, 2003. 22(5): p. 421–441.

39. Browne, R.K., et al., Sperm motility of externally fertilizing fish and amphibians. Theriogenology, 2015. 83(1): p. 1-13.e8.

40. Spindler, R., T. Keeley, and N. Satake, Applied andrology in endangered, exotic and wildlife species, in Animal andrology: theories and applications. 2014, CABI Wallingford UK. p. 450–473.

41. van der Horst, G., et al., Testicular Structure and Spermatogenesis in the Naked Mole-Rat Is Unique (Degenerate) and Atypical Compared to Other. Sperm Differentiation and Spermatozoa Function: Mechanisms, Diagnostics, and Treatment, 2020: p. 230089.

42. Van der Horst, G., et al., Testicular structure and spermatogenesis in the naked mole-rat is unique (degenerate) and atypical compared to other mammals. Frontiers in Cell and Developmental Biology, 2019. 7: p. 234.

43. Chen, L., et al., Electroejaculation and semen characteristics of Asiatic black bears (Ursus thibetanus). Animal reproduction science, 2007. 101(3-4): p. 358–364.

44. Morato, R., et al., Comparative analyses of semen and endocrine characteristics of free-living versus captive jaguars (Panthera onca). Reproduction, 2001. 122(5): p. 745–751.

45. Gañán, N., et al., Assessment of semen quality, sperm cryopreservation and heterologous IVF in the critically endangered Iberian lynx (Lynx pardinus). Reproduction, Fertility and Development, 2009. 21(7): p. 848–859.

46. Terrell, K.A., et al., Evidence for compromised metabolic function and limited glucose uptake in spermatozoa from the teratospermic domestic cat (Felis catus) and cheetah (Acinonyx jubatus). (1529-7268 (Electronic)).

47. Martínez-Torres, M., et al., Semen Evaluation from Dominant Males of the Viviparous Mexican Lizard Sceloporus torquatus, Wiegmann, 1828 (Sauria: Phrynosomatidae). Veterinary Sciences, 2025. 12(4): p. 363.

48. Simmons, L.W. and N. Wedell, Fifty years of sperm competition: The structure of a scientific revolution. 2020, The Royal Society. p. 20200060.

49. Browne, R.K., et al., Sperm motility of externally fertilizing fish and amphibians. Theriogenology, 2015. 83(1): p. 1–13.

50. Brandies, P., et al., The Value of Reference Genomes in the Conservation of Threatened Species. Genes, 2019. 10(11): p. 846.

51. Bertola, L.D., et al., A pragmatic approach for integrating molecular tools into biodiversity conservation. Conservation Science and Practice, 2024. 6(1): p. e13053.

52. Byers, B. and A. Cameron, INTEGRATING CLIMATE CHANGE ADAPTATION INTO BIODIVERSITY AND FORESTRY ASSESSMENTS AND PROGRAMMING Integrating Climate Change Adaptation into Biodiversity and Forestry Assessments and Programs PREPARED BY TABLE OF CONTENTS Abbreviations and Acronyms ………………………………………………………………………..i Acknowledgements …………………………………………………………………………………….iii. 2013.

53. Holderegger, R., et al., Conservation genetics: Linking science with practice. Molecular Ecology, 2019. 28(17): p. 3848–3856.

54. Waggener, W.L. and E.J. Carroll Jr, Spermatozoon structure and motility in the anuran Lepidobatrachus laevis. Development, growth & differentiation, 1998. 40(1): p. 27–34.

55. Waggener, W.L. and E.J. Carroll Jr, A method for hormonal induction of sperm release in anurans (eight species) and in vitro fertilization in Lepidobatrachus species. Development, growth & differentiation, 1998. 40(1): p. 19–25.

56. Christensen, J.R., et al., Validation of an amphibian sperm inhibition toxicological test method using zinc. Environmental toxicology and chemistry, 2004. 23(12): p. 2950–2955.

57. Pearl, E., et al., An optimized method for cryogenic storage of Xenopus sperm to maximise the effectiveness of research using genetically altered frogs. Theriogenology, 2017. 92: p. 149–155.

58. Ueda, Y., Y. Yoshizaki N Fau - Iwao, and Y. Iwao, Acrosome reaction in sperm of the frog, Xenopus laevis: its detection and induction by oviductal pars recta secretion. (0012-1606 (Print)).

59. Arregui, L., et al., Effect of seasonality on hormonally induced sperm in Epidalea calamita (Amphibia, Anura, Bufonidae) and its refrigerated and cryopreservated storage. Aquaculture, 2020. 529: p. 735677.

60. Kouba, A.J., et al., Structural and functional aspects of Bufo americanus spermatozoa: effects of inactivation and reactivation. Journal of Experimental Zoology Part A: Comparative Experimental Biology, 2003. 295(2): p. 172–182.

61. Guy, E.L., et al., Evaluation of different temporal periods between hormone-induced ovulation attempts in the female Fowler’s toad Anaxyrus fowleri. Conservation Physiology, 2020. 8(1): p. coz113.

62. Browne, R.K., et al., Sperm collection and storage for the sustainable management of amphibian biodiversity. Theriogenology, 2019. 133: p. 187–200.

63. Silla, A.J., A.J. Kouba, and H. Heatwole, Reproductive technologies and biobanking for the conservation of amphibians. 2022: CSIRO PUBLISHING.

64. Chen, D.M., et al., The impact of time and environmental factors on the mitochondrial vesicle and subsequent motility of amphibian sperm. Comparative Biochemistry and Physiology Part A: Molecular & Integrative Physiology, 2022. 268: p. 111191.

